# Ascites-Driven Modulation of Cell Phenotypes and Proteomes: Implications for Cancer Progression

**DOI:** 10.64898/2026.02.12.705674

**Authors:** Jack Scanlan, Parul Mittal, Jennifer Washington, Clifford Young, Martin K Oehler, Peter Hoffmann, Manuela Klingler-Hoffmann

## Abstract

**Background:** More than 90% of advanced ovarian cancer patients develop malignant ascites, which describes a buildup of fluid in the peritoneal cavity caused by increased vascular permeability and obstructed lymphatic drainage. Malignant ascites contains spheroidal tumor cell clusters that contain stromal cells, cancer-associated fibroblasts, and blood cells. These spheroids promote peritoneal metastasis and treatment resistance, yet the phenotypic and proteomic changes of cells caused by the ascites environment remain poorly understood, as does its influence on *ex vivo* responses to chemotherapeutics in personalized medicine approaches.

**Methods:** Using mass spectrometry, we compared the proteome profiles of cell-free ascites to serum from ovarian cancer patients. We then analyzed the proteomes of immortalized cancer cells grown as monolayers or spheroids in either malignant ascites or standard cell culture medium. The effects of this fluid on the phenotype, molecular composition, and *ex vivo* chemotherapy responses of cancer cells were also investigated.

**Results:** Proteome analysis revealed that cell-free ascites had higher levels of extracellular, secreted, and membrane proteins compared to serum. Ascites enhanced cell viability and spheroid formation in immortalized ovarian cancer cell lines more effectively than standard cell culture medium. Despite this altered baseline viability, growth of spheroids in ascites versus cell culture medium did not hinder chemotherapy response assessments, indicating the appropriateness of standard cell culture medium in *ex vivo* applications. The observed phenotypic changes of cells grown in ascites could not be recapitulated by adding chemokines or periostin to the cell culture medium, suggesting that additional factors are required. Notably, elevated levels of transglutaminase 2 were identified in SKOV-3 cells grown in ascites, indicating that ascites directly influences protein expression in cancer cells.

## 1. Introduction

Ovarian cancer remains one of the most lethal gynecological malignancies, which is primarily attributed to advanced-stage diagnosis and innate or acquired chemotherapy resistance. Despite advances in surgical techniques and chemotherapy regimens, the five-year survival rate of advanced ovarian cancer patients has remained below 50 per cent for several decades [1]. Ovarian cancer often correlates with the presence of ascites— the accumulation of fluid within the peritoneal cavity. While most patients with advanced disease accumulate more than 2 liters of ascites [2], even a small volume of ≥ 100 mL can indicate poor prognosis [3]. This buildup of fluid contributes to patient discomfort and morbidity and creates a complex tumor microenvironment due to the presence of aggregated cells, known as spheroids, that shed from the primary tumor [4]. Malignant ascites can also contain immune cells, extracellular vesicles [5-7], cytokines, growth factors, enzymes, and cell-free DNA [8], which have been shown to contribute to cell survival, proliferation, and dissemination of cancer cells [9, 10] enabled through direct contact with the peritoneal lining [11].

The conserved genetic landscape of ascitic tumor material compared to primary ovarian tumors [12], as well as their influence on chemoresistance and metastasis, make patient-derived ascites an attractive tool with which to personalize treatment. Retrospective *ex vivo* treatment response assessments on three-dimensional ascitic cell cultures have recapitulated patients’ clinical responses to platinum-based chemotherapy [13-18], paclitaxel and inhibitors of PARP [14], and monoclonal antibody treatments [17]. Furthermore, prospective studies have used such data to successfully guide treatment selection [19, 20]. However, few studies analyze their responses to different *ex vivo* culture conditions and their effect on the validity of *ex vivo* chemoresponse assessments. This is especially important given that alterations in growth conditions of ascites cells, differentially regulated cultures versus spheroidal cultures, have enabled the identification of a number of differentially-regulated proteins such as the cell cycle arrest marker Aurora kinase B [21].

Proteomic and metabolomic profiling techniques have expanded our view of the ascitic milieu and highlight its role in disease progression. For instance, altered T-cell function, interleukin pathway upregulation, and reduced MHC-II expression reduce anti-tumor and immunosuppressive effects to promote disease progression [22-24]. Additional factors within ascites can also promote metastasis; for instance, through TGF-β1-mediated increases to the proliferation and migration of mesothelial cells [25]. Comparative studies of cirrhotic and malignant ascites have also identified cancer-specific pathways, metabolites, and spliceosomal components [26], further demonstrating the role of microenvironment tumor biology.

The identification of altered pathways between chemotherapy-treated and chemotherapy-naïve ovarian cancer patients [27] demonstrates the utility of proteomic analyses in personalized medicine approaches. For instance, researchers identified a 64-protein signature that is capable of reliably forecasting whether high-grade serous ovarian tumors would respond to standard therapies with 98 percent specificity [28]. While more limited, some studies have recognized the value of ascites as a window into the evolving molecular landscape of individual ovarian cancers, enhancing the detection of candidate biomarkers that could be pivotal in early diagnosis and the personalization of treatments. For example, an early study identified ceruloplasmin by gel-based MALDI-TOF/TOF mass spectrometry as a marker of treatment resistance in cell-free ascites and validated this finding by ELISA in 28 patients [29]. Using advanced data-independent acquisition-based mass spectrometry, our laboratory recently demonstrated that proteomic profiling can distinguish the clinical treatment responses of patients’ ascitic spheroids [30]. However, the poor alignment of analytical techniques, sample types, treatment regimens, and reporting standards between such studies [31] have hindered the generalizability of findings.

Despite evidence of malignant ascites as a useful biofluid for ovarian cancer, our understanding of its composition and utility in *ex vivo* chemoresponse testing, as well as its impacts on spheroid formation, remains limited. Here, we employ morphological and proteomics approaches to explore how the ascitic microenvironment alters spheroid aggregation and protein expression. The complexity of this biofluid was first compared to patient-derived serum, with distinct differences in the identified ECM, secreted, and membrane proteins between both serum and ascites, and between ascites sourced from different patients with vastly different spheroid-forming capabilities. This translated into distinct morphological effects of ascites on immortalized ovarian cancer cell lines. We demonstrate that barrier-forming cornification and keratinization pathways are significantly enriched in non-spheroid-promoting ascites, compared to collagens and laminins in spheroid-promoting ascites. We also used adherent and spheroidal cultures to demonstrate that ascites promotes the upregulation of spheroid-promoting protein TGM2. In the context of ascites cells as a tool for *ex vivo* treatment response assessments, we show that the ascitic environment only confers minor survival protection that does not significantly alter the cells’ chemotherapy responses compared to standard cell culture medium. These data pave the way for simplified *ex vivo* treatment response assessments that could predict patients’ responses to chemotherapy before administration.

## 2. Results

### 2.1. Comparative proteomics of depleted ascites

Ascites from three ovarian cancer patients, named Ascites 1, 2, and 3, were collected during cytoreductive surgery prior to adjuvant chemotherapy. The volume of ascites, number of cells, and presence of spheroids within the malignant ascites varied between patients. To identify proteins that are specifically enriched in the tumor microenvironment from those commonly found in circulation, we compared the proteomes of ascites from the 3 patients to healthy serum. Since serum contains a high concentration of abundant proteins such as albumin and immunoglobulins which can interfere with the identification of the low-abundant proteins, the top-14-most abundant proteins were depleted prior to mass spectrometric analysis in triplicate or quadruplicate. Of the 845 proteins identified across all runs, 793 remained after removal of potential contaminants, reverse proteins, and proteins only detected at the MS1 level. Of these, 298 were detected in all 4 samples (Figure 1a) and 67 were uniquely identified in ascites from all patients, suggesting potential tumor microenvironment-specific proteins. Interestingly, 111 proteins were detected only in the serum but not in the ascites samples. Several proteins were found to be specific to individual ascites samples, including 63 unique to Ascites 1, 95 to Ascites 2, and 54 to Ascites 3, reflecting patient-specific variability.

**Figure 1.**
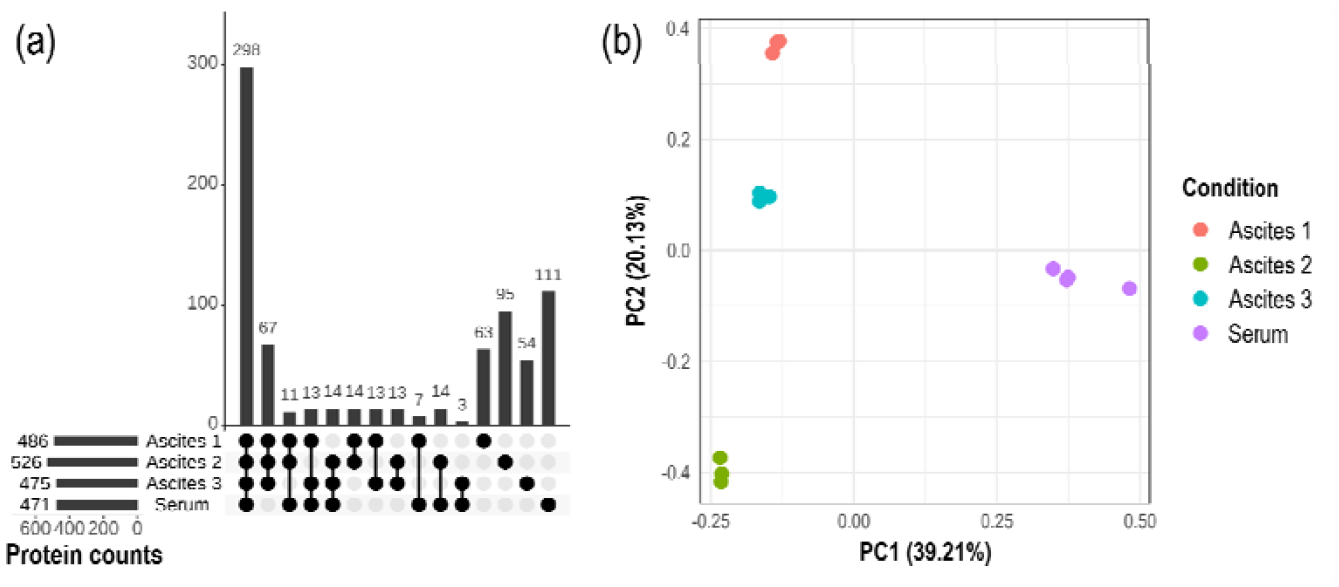
Comparative proteomic analysis of serum and ascites from three patients identified proteins specific to ascites. (a) An upset plot showing shared and unique protein groups across three depleted ascites samples and depleted serum, generated with the UpsetR package [32]. (b) A principal component analysis separated ascites from serum along the first component, and ascites from different patients along the second component. Low variability was observed between technical replicates.

Importantly, a clear distinction was observed between the serum and ascites samples along the first component of the PCA plot, with ascites samples separated by patient origin along the second component (Figure 1b). Technical replicates resulted in highly reproducible data with minor technical variability.

To identify proteins enriched in the ascitic tumor microenvironment, we focused on secreted and extracellular matrix (ECM) proteins that were detected in all ascites samples but not in serum. Of the 67 proteins detected in all ascites samples but absent from serum, 19 were either secreted, ECM, or membrane proteins (Table 1).

**Table 1.**
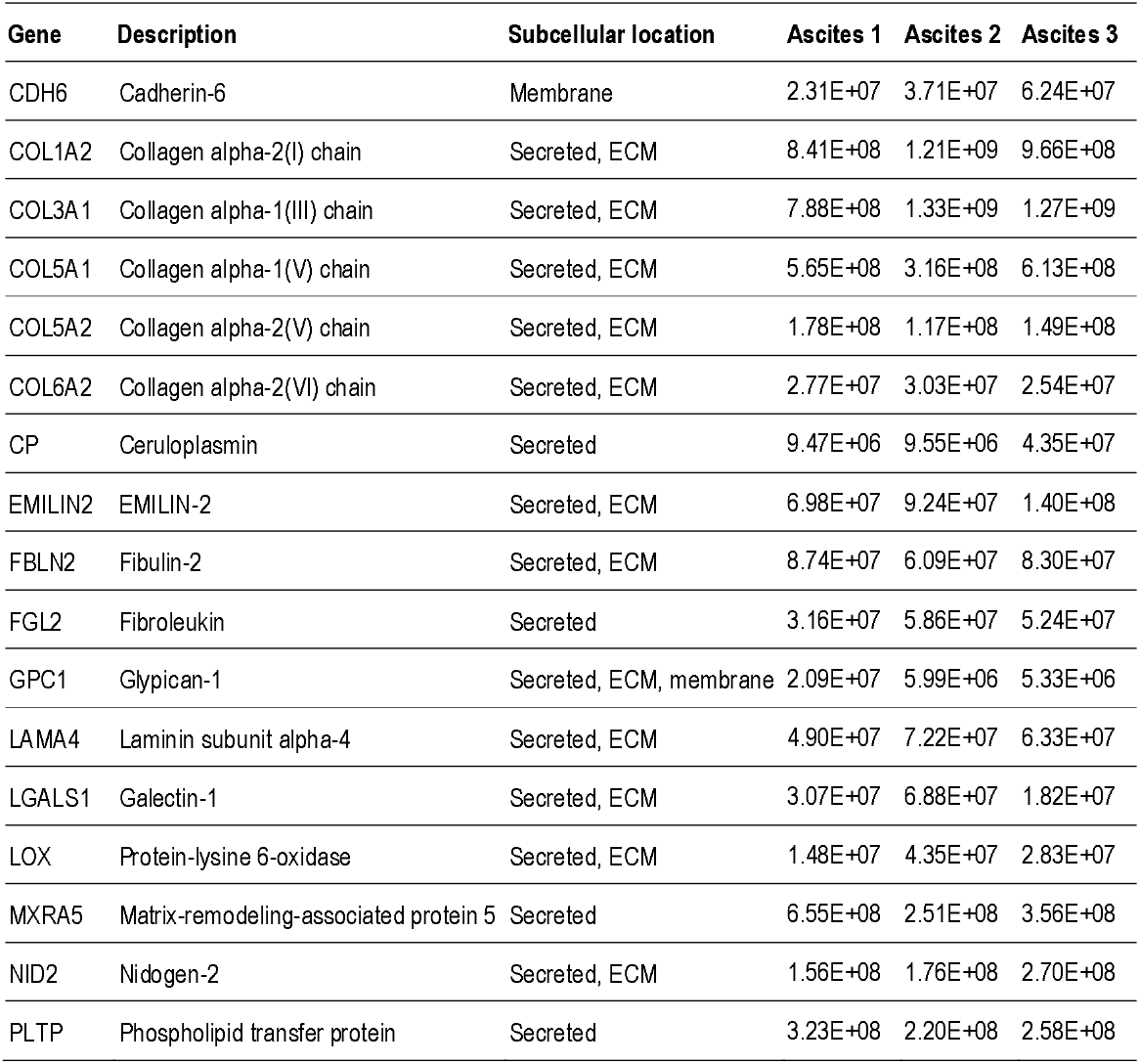
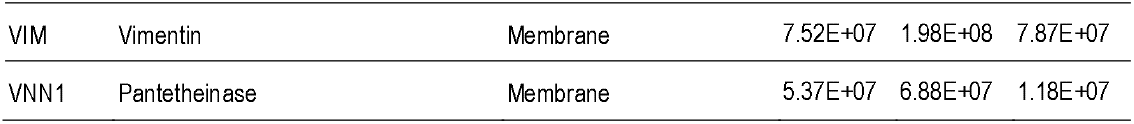
Average abundances of secreted, ECM, or membrane proteins, which are absent in serum but detected in all ascites. Average protein abundance calculated from triplicate measurements for each sample, based on LFQ intensity.

| Gene | Description | Subcellular location | Ascites 1 | Ascites 2 | Ascites 3 |
| --- | --- | --- | --- | --- | --- |
| CDH6 | Cadherin-6 | Membrane | 2.31E+07 | 3.71E+07 | 6.24E+07 |
| COL1A2 | Collagen alpha-2(I) chain | Secreted, ECM | 8.41E+08 | 1.21E+09 | 9.66E+08 |
| COL3A1 | Collagen alpha-1(III) chain | Secreted, ECM | 7.88E+08 | 1.33E+09 | 1.27E+09 |
| COL5A1 | Collagen alpha-1(V) chain | Secreted, ECM | 5.65E+08 | 3.16E+08 | 6.13E+08 |
| COL5A2 | Collagen alpha-2(V) chain | Secreted, ECM | 1.78E+08 | 1.17E+08 | 1.49E+08 |
| COL6A2 | Collagen alpha-2(VI) chain | Secreted, ECM | 2.77E+07 | 3.03E+07 | 2.54E+07 |
| CP | Ceruloplasmin | Secreted | 9.47E+06 | 9.55E+06 | 4.35E+07 |
| EMILIN2 | EMILIN-2 | Secreted, ECM | 6.98E+07 | 9.24E+07 | 1.40E+08 |
| FBLN2 | Fibulin-2 | Secreted, ECM | 8.74E+07 | 6.09E+07 | 8.30E+07 |
| FGL2 | Fibroleukin | Secreted | 3.16E+07 | 5.86E+07 | 5.24E+07 |
| GPC1 | Glypican-1 | Secreted, ECM, membrane | 2.09E+07 | 5.99E+06 | 5.33E+06 |
| LAMA4 | Laminin subunit alpha-4 | Secreted, ECM | 4.90E+07 | 7.22E+07 | 6.33E+07 |
| LGALS1 | Galectin-1 | Secreted, ECM | 3.07E+07 | 6.88E+07 | 1.82E+07 |
| LOX | Protein-lysine 6-oxidase | Secreted, ECM | 1.48E+07 | 4.35E+07 | 2.83E+07 |
| MXRA5 | Matrix-remodeling-associated protein 5 | Secreted | 6.55E+08 | 2.51E+08 | 3.56E+08 |
| NID2 | Nidogen-2 | Secreted, ECM | 1.56E+08 | 1.76E+08 | 2.70E+08 |
| PLTP | Phospholipid transfer protein | Secreted | 3.23E+08 | 2.20E+08 | 2.58E+08 |
| VIM | Vimentin | Membrane | 7.52E+07 | 1.98E+08 | 7.87E+07 |
| VNN1 | Pantetheinase | Membrane | 5.37E+07 | 6.88E+07 | 1.18E+07 |

### 2.2. Phenotype of cells grown as three-dimensional cell clusters in ascites and medium

#### 2.2.1. Impact of patient-derived ascites on 3D ovarian cancer cell growth

To investigate the effect of ascites on primary tumor cell aggregation and cell cluster morphology, primary ovarian cancer cells were cultured in ultra-low attachment plates as 2D spheroids with increasing concentrations of ascites (Supplementary Figure S1). Spheroid size and compactness increased with higher concentrations of malignant ascites, with the most pronounced effect observed in ascites compared to medium alone (data not shown). In addition, adherent ascites cells from two patients that were cultured in either standard medium or patient-matched ascites formed larger and more compact spheroids than those grown in standard medium (representative images in Figure 2a).

**Figure 2:**
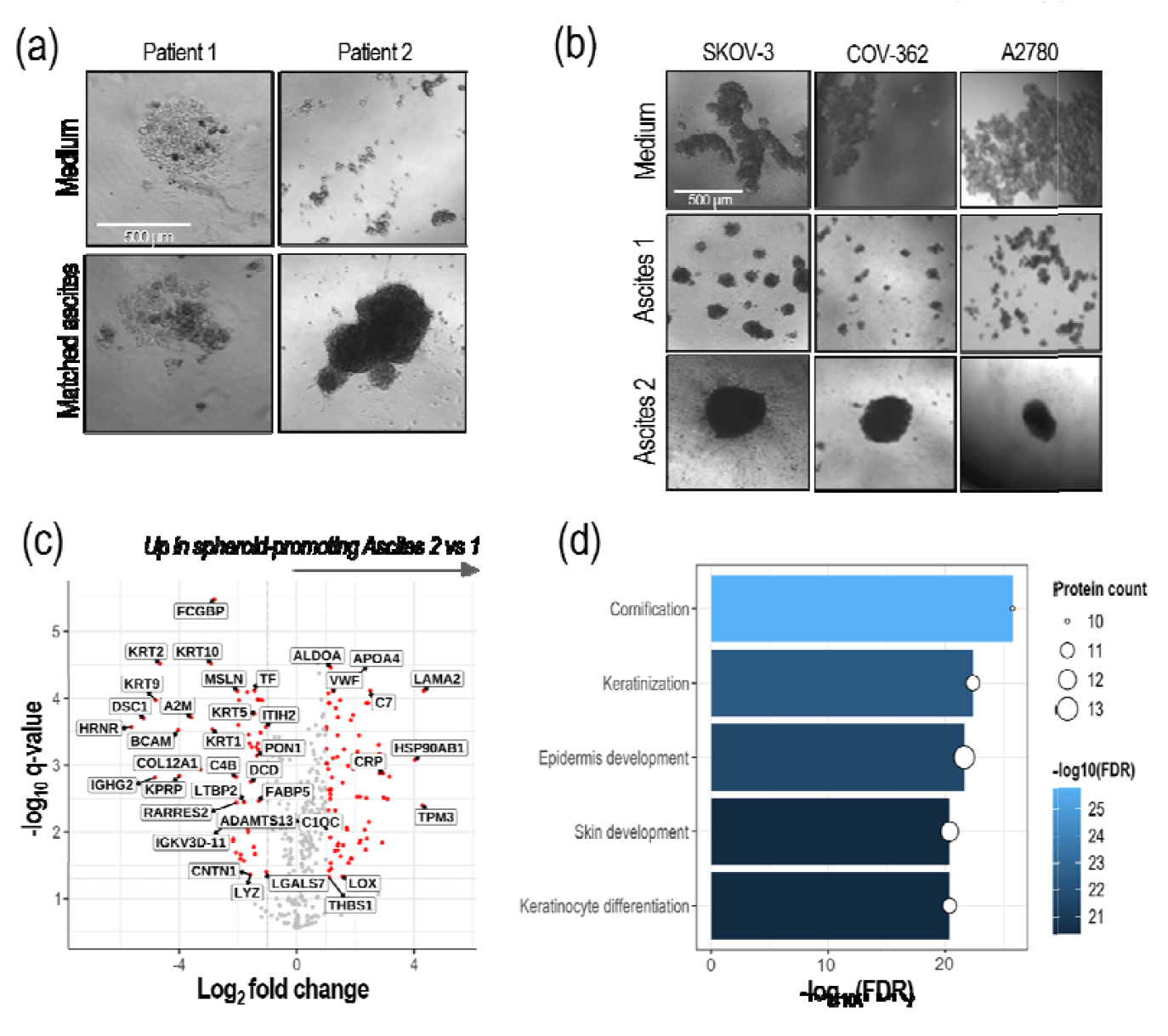
Ascites induces phenotypic changes in three-dimensional cell clusters. (a) Ascites was collected from patients and spun down. 2000 cells were plated per well in 96-well ultra-low adhesive plates, cultured in medium or 100% ascites and imaged after 72 hours using an inverted microscope. Row 1 shows cells from patient 1 grown as spheroids in medium or ascites 1 and row 2 than those incubated in standard cell culture medium However the degree of cell death

To explore whether a similar effect could be observed with ovarian cancer cell ines, we repeated the experiment with SKOV-3, COV362, and A2780 cells. All cell lines formed more compact and organized spheroids in ascites compared to medium, suggesting that components in ascites promote aggregation (Figure 2b). However, cells grown in Ascites 2 formed larger and tighter spheroids than those grown in Ascites 1, indicating that molecular differences in ascites can alter three-dimensional ovarian cancer cell phenotypes. and A2780 cells were cultured per well in DMEM or 100% ascites and imaged after 72 hours cells using an inverted microscope. All images are one representative example of three technical replicates. (c) The fold-change and significance of proteins observed in spheroid-promoting Ascites 2 versus Ascites 1 displayed in a volcano plot created with EnhancedVolcano R package [32], with (d) biological processes enriched in proteins uniquely detected in Ascites 1 versus Ascites 2 analyzed using DAVID (NIH) [33].

We explored the proteomic differences between Ascites 1 and 2 to discover which proteins could contribute to the observed phenotypical changes. The proteomics data were analysed using a volcano plot (Figure 2c). Basal cell adhesion molecule (BCAM), hornerin (HRNR), and keratins (KRT1/KRT2/KRT5/KRT9/KRT10) were among those more abundant in Ascites 1, whereas heat shock protein 90 isoforms (HSP90AB1, HSP90B1, HSP90AA1), LAMA2, S100A8/S100A9, and insulin growth factor 2 (IGF2) were among the proteins more abundant in Ascites 2. Kallikreins (KLK5/KLK6/KLK10) and other barrier-forming proteins were uniquely present in Ascites 1, whereas periostin (POSTN), growth factors (CTGF, EGFR) and integrins (ITGB2, ITGAM) were uniquely present in Ascites 2. Resulting enriched biological processes are summarised in Figure 2d.

#### 2.2.2 Neither cytokines nor periostin induce significant phenotypic changes in 3D-cultured ovarian cancer cells

Given the unique presence of POSTN in spheroid-promoting Ascites 2, we assessed whether its inclusion in cell culture medium can induce a phenotypic change during SKOV-3 spheroid generation. This was expanded to a panel of selected pro-inflammatory and tumor-associated cytokines given their well-documented roles in modulating cell proliferation, survival, inflammation, and ovarian cancer progression [33]. After 72 and 144 hours, SKOV-3 cells formed spheroids under all conditions, with moderate variation in spheroid sizes and shapes (Figure 3). These data indicate that neither the addition of cytokines nor POSTN, neither alone nor in combination, were sufficient to induce a change in phenotype as it has been observed when the cells were seeded in Ascites 2. Neither condition altered viability (data not shown).

**Figure 3:**
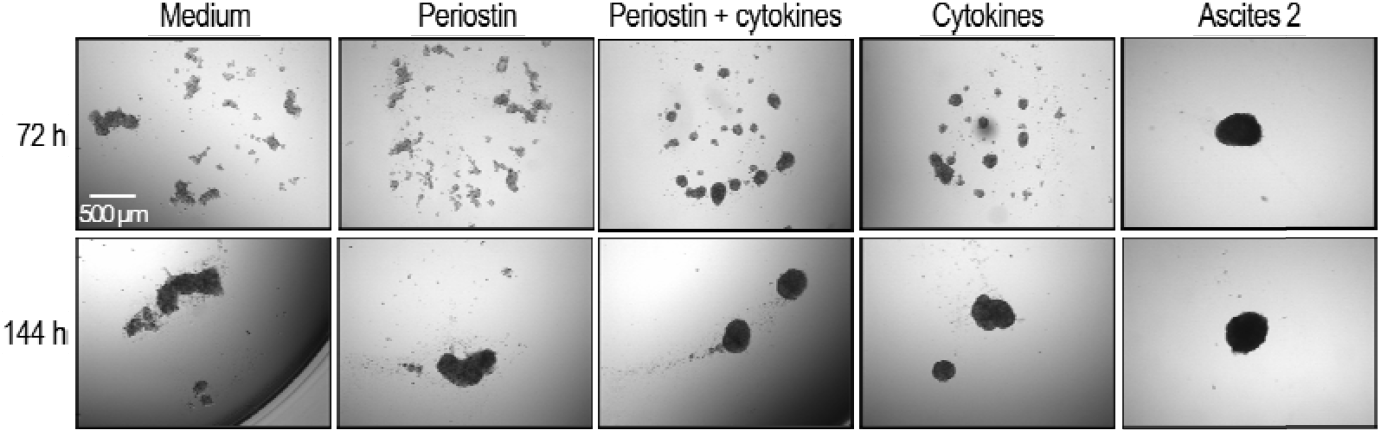
Cytokines and POSTN induce minimal phenotypic changes of SKOV-3 cells grown in 3D. SKOV-3 cells were grown in medium, medium supplemented with cytokines (0.25 ng/ml IL1-β, 1 ng/ml IL6, 10 ng/ml TNFα, 10 ng/ml TGFβ, medium supplemented with 50 ng/well POSTN or 100% Ascites 2. After 72 hours and 144 hours the cells were imaged using an inverted microscope. Shown is one representative well of cells seeded in triplicates in a 96-well plate.

To investigate whether the phenotypic changes induced by ascites changes chemotherapeutic response, SKOV-3 cells were cultured in 3D using patient-derived Ascites 1 and Ascites 2. More compact and larger spheroids formed in the presence of ascites, particularly with Ascites 2, and the viability of cells cultured in ascites was higher than those incubated in standard cell culture medium However the degree of cell death following carboplatin treatment did not differ, suggesting that phenotypic changes alone are not predictive of drug response (data not shown).

#### 2.2.3 Proteomic analysis of SKOV-3 cells grown in patient ascites

Given the lack of effect of interleukins and POSTN on three-dimensional immortalized cell cultures, we aimed to understand how ascites-associated factors affect ovarian cancer cells. To do this, SKOV-3 cells were grown as spheroids in standard DMEM medium alone, or Ascites 1 and Ascites 2. We then performed a detailed proteomic analysis to compare protein expression in SKOV-3 cells grown under these different conditions. Each sample was analyzed in triplicate to ensure consistent and reliable results.

Of the 6,142 proteins detected across all samples, 6074 remained following the removal of potential contaminants, reverse, and proteins detected only at MS1 level (Figure 4a). Of these, 3234 were identified in all conditions, with 968 proteins uniquely iden ified in 3D SKOV-3 cells grown in ascites of any patient origin and 165 commonly identified in spheroids cultured in both ascites. Interestingly, 1174 proteins were uniquely iden ified in spheroids grown in standard medium, with this difference reflected in the segregation of all samples in the principal component analysis (Figure 4b). However, hierarchical clustering grouped spheroids that were cultured in both ascites separately from those grown in medium alone (Supplementary Figure S2). These data show that ascites significantly changes the proteome of ovarian cancer cells, highlighting how the tumor microenvironment can influence cancer cell biology and possibly affect tumor growth and response to therapy.

**Figure 4.**
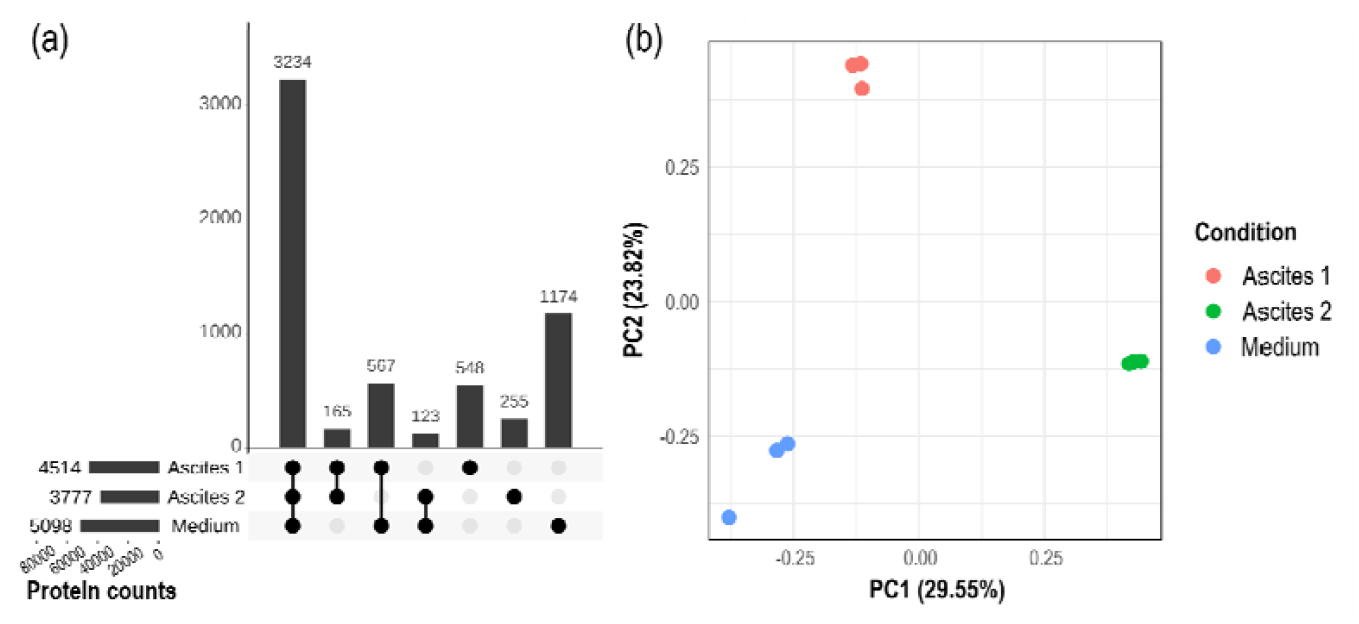
Comparative proteomic analysis of SKOV-3 cells grown as 3D tumor spheroids in DMEM Medium, Ascites 1 or Ascites 2. The intersection of all quantified proteins shown as an upset plot generated with the UpsetR package [32] with (b) PCA plot showing separation of each group.

The abundances of secreted, ECM, and membrane proteins in SKOV-3 spheroids were compared between those grown in medium and those grown in ascites from two separate patients, and are listed in Supplementary Table S1. Collagens (1A1, 1A2, 3A1, 5A1, 5A2, 6A2), apolipoproteins (A-I, A-IV, C-IV, D, E), were significantly more abundant in spheroids cultured in patient ascites, with many not detected in samples cultured in standard cell culture medium.

Coagulation factors (X, XI, XII), haptoglobin, hemopexin, alpha-1B-glycoprotein, and ceruloplasmin are plasma proteins that were detected only in ascites-cultured spheroids. Given that malignant ascites originates from increased capillary permeability, such proteins may simply be derived from the ascites, rather than produced by spheroids in response to their environment. To clarify, we repeated the experiment using adherent cultures that can be processed more vigorously to reduce contamination of proteins, which may stick to surfaces and cells. In this setup, SKOV-3 cells were grown in monolayer using either standard DMEM medium or 100% ascites from two patients. Culturing the cells adherently allowed for more rigorous washing steps to remove residual medium or ascites before protein extraction, thereby minimizing potential contamination from external proteins (Figure 5a).

**Figure 5.**
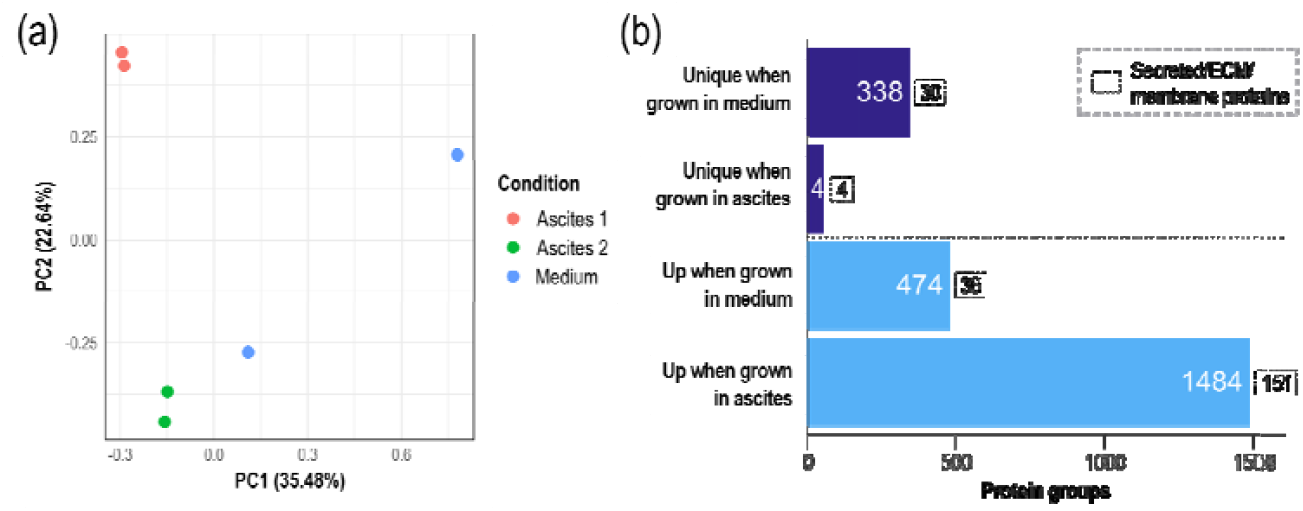
Comparative proteomic analysis of adherent 2D SKOV-3 cells cultured in standard cell culture medium, or ascites from two patients. (a) PCA plot summarizing protein abundance information from duplicate mass spectrometer injections of cells grown in each culture condition. (b) Uniquely present and upregulated proteins in adherent SKOV-3 cells that were cultured in medium and malignant ascites, annotated with the number which are classified as secreted, or reside in the ECM or membrane.

The number of upregulated proteins in adherent SKOV-3 cells that were cultured in medium and ascites were assessed (Figure 5b). From 812 proteins that were elevated in medium-cultured 2D SKOV-3 cells, including several cell cycle regulation proteins, approximately half were exclusive to this condition. In contrast, less than 5% of proteins that had higher abundances in ascites-cultured 2D SKOV-3 cells were detected in both culture conditions, with most instead being identified exclusively in cells cultured with patient ascites.

Given their roles in spheroid formation, we also assessed the number of secreted, ECM, and membrane proteins that were upregulated in either culture condition. The vast majority (155) were upregulated or uniquely present in cells grown in ascites, with 151 being uniquely identified in this culture condition. Of these proteins, 22 were also more abundant in 3D cultures compared to those cultured in standard medium (Table 3). Twenty-one were only identified in adherent cells cultured in ascites, with apolipoproteins, collagens, coagulation factors, and cadherin 6 commonly detected in 2D and 3D ascites cultures. Only TGM2, was present in both medium and ascites-cultured spheroidal and adherent cultures while being elevated in ascites-cultured cells.

**Table 3.** Alphabetized secreted, ECM, and membrane proteins that were upregulated in both two- and three-dimensional SKOV-3 cells cultured in all ascites compared to those cultured in medium, with the average relative abundances of two-dimensional cultures from all technical replicates of those cultured in ascites (n=2) and medium presented. TGM2 is highlighted as the only listed protein that was identified in ascites and medium culture conditions. (ND=not detected).

| Gene | Description | Subcellular location | Ascites | Medium |
| --- | --- | --- | --- | --- |
| A1BG | Alpha-1B-glycoprotein | Secreted | 5.23E+08 | ND |
| APOA1 | Apolipoprotein A-I | Secreted | 2.23E+10 | ND |
| APOA4 | Apolipoprotein A-IV | Secreted | 1.26E+09 | ND |
| APOB | Apolipoprotein B-100 | Secreted | 1.03E+10 | ND |
| APOD | Apolipoprotein D | Secreted | 4.46E+08 | ND |
| APOE | Apolipoprotein E | Secreted, ECM | 5.00E+09 | ND |
| CDH6 | Cadherin-6 | Membrane | 2.39E+06 | ND |
| COL3A1 | Collagen alpha-1(III) chain | Secreted, ECM | 2.29E+07 | ND |
| COL5A1 | Collagen alpha-1(V) chain | Secreted, ECM | 3.64E+07 | ND |
| COL5A2 | Collagen alpha-2(V) chain | Secreted, ECM | 2.14E+07 | ND |
| CP | Ceruloplasmin | Secreted | 2.21E+09 | ND |
| CST3 | Cystatin-C | Secreted | 1.93E+08 | ND |
| F11 | Coagulation factor XII | Secreted | 1.13E+08 | ND |
| F12 | Coagulation factor XII | Secreted | 2.09E+07 | ND |
| F2 | Prothrombin | Secreted, ECM | 6.26E+08 | ND |
| FGB | Fibrinogen beta chain | Secreted | 1.89E+09 | ND |
| GC | Vitamin D-binding protein | Secreted | 1.81E+09 | ND |
| HP | Haptoglobin | Secreted | 3.20E+10 | ND |
| HPX | Hemopexin | Secreted | 9.62E+08 | ND |
| PLG | Plasminogen | Secreted | 8.20E+08 | ND |
| **TGM2** | **Protein-glutamine**<br>**gamma-glutamyltransferase 2** | **Secreted, ECM, membrane** | **1.73E+09** | **6.22E+08** |
| VTN | Vitronectin | Secreted, ECM | 2.30E+09 | ND |

## 3. Discussion

Many ovarian cancer cases present with malignant ascites containing tumor cells, yet the interplay between ascites and the aggregation of these cells into spheroids remains unclear. First, we compared the proteomic profiles of ascites from three patients to serum, Importantly, 67 proteins were uniquely present in all ascites samples, with 19 of these being secreted, located in the ECM and/or the cell membrane.

The ECM consists of more than 300 proteins, which not only provide structural stability, but also regulate key cellular processes that drive cancer development [29]. Collagens are of particular interest in cancer cell survival, with several being identified as molecular markers of disease progression and chemoresistance through multi-omics approaches. For instance, the degradation of collagen 1 into glutamine for resorption by tumor cells has been shown to satisfy their high metabolic demand [34]. Collagen density, organization, and signaling also promote metastasis [35] and chemotherapy resistance [36], potentially through increased ECM stiffness that modulates drug efflux [37]. Importantly for ovarian cancer progression, increased ECM density promotes larger spheroid clusters [38]. This has led to the development of commercial products that are designed to drive the aggregation and formation of spheroids. Our data show that several collagens are uniquely present in all ascites samples, alongside other ECM proteins also implicated in disease progression, such as cadherin-6 [39], laminin subunit alpha-4 [40], and fibulin 2 [41]; however, their complex and often contrasting roles have prevented their use as molecular markers of disease.

Although not classical ECM proteins, the identification of several ECM-regulating annexins (ANXA1, ANXA2, ANXA5) solely in ascites supports the proposed function of this protein family as molecular markers of ovarian cancer. For instance, ANXA2 overexpression has been observed in ovarian cancer tissues, and its upregulation has been linked to worsened overall survival and ascites. However, the direction of change is often inconsistent between studies, with proteomic studies identifying annexins as markers of cisplatin [42] and paclitaxel [43] resistance. Other proteins uniquely detected in ascites samples include laminins and SPON1, both of which have been implicated in disease progression and metastasis [44, 45]. Importantly, the present study demonstrates that ascites may be a more suitable biofluid for the detection of ECM and secreted proteins and their known regulators.

The morphologies of cell line spheroids that were cultured in ascites sourced from different patients varied, which was replicated in patient-derived primary ascites cells. This supports the distinct effects of ascites fluid from different patients, potentially resulting from the differing proteome profiles of ascites. This was demonstrated in the significant upregulation of several heat shock 90 isoforms (e.g., HSP90AB1, HSP90B1, and HSP90AA1) in the spheroid-promoting Ascites 2, along with S100A8/S100A9 [46] and integrins (e.g., ITGB1) [47], which have been reported as potential negative anoikic regulators. Several relevant proteins were also uniquely present in spheroid-promoting Ascites 2 but not Ascites 1, including POSTN, growth factors (CTGF, EGFR) and integrins (ITGB2, ITGAM). CTGF is a well-established downstream mediator of TGF-β that drives COL1A1 and broader collagen I deposition in fibrotic and tumor settings, which has been shown to support tightly compacted, drug-resistant tumor spheroids [48]. These data suggest that Ascites 2 provides a CTGF-, integrin-and EGFR-competent ECM that favors collagen I–dependent cell–to-cell and cell–to-matrix adhesion, consistent with the more compact spheroid morphologies observed in both cell lines and primary ascites cells. A large subset of proteins only identified in Ascites 1 instead promote barrier and cornified envelope formation (e.g., FLG/FLG2, KLK5/KLK6/KLK10, and XP32), which typically stiffens stratified epithelia [49]. In this context, such proteins are likely to reduce motility and spheroid formation.

Given the key differences in secreted and ECM proteins present within ascites and standard cell culture medium, the distinct proteomic profiles and morphologies of spheroids grown in different environments raise questions regarding the validity of *ex vivo* dose response assessments performed in standard cell culture medium. Our observation of a reduced baseline viability when SKOV-3 spheroids are grown in standard medium rather than patient-derived ascites fluid supports previous findings [50]; however, we report that standard cell culture media supports robust dose response testing with CBP, the most common platinum-based ovarian cancer chemotherapeutic.

We have previously observed that the sizes of spheroids generated from both cell lines and patient-derived ascites cells do not correlate with their *ex vivo* chemoresponses, nor the clinical outcomes of patients [29]. Given the role of spheroids as metastatic units that contribute to cancer dissemination and treatment resistance, future studies should assess whether the spheroid-forming capability of ascites can correlate with patient survival and chemoresponse. Nevertheless, we hypothesized a bidirectional relationship between the patient-dependent aggregation capabilities of ascites alongside standard cell culture medium and the proteomes of cultured spheroids. We used mass spectrometry to profile the proteomes of SKOV-3 cells that were cultured as three-dimensional spheroids in ascites versus cell culture medium, which emulates the micro-environment present in malignant ascites. Secreted and ECM proteins produced by tumor or stromal cells mediate cell-to-cell communication, tumor growth, and metastasis [51], therefore were the focus of our analyses. The abundance of apolipoproteins in ascites-cultured SKOV-3 spheroids, which regulate lipid metabolism, transport, and enzyme activity in cancer [52] reflect a complex microenvironment. The elevated or unique abundance of collagens in ascites-cultured spheroids reflects their previously discussed reported roles in spheroid formation and chemoresponse.

A previous proteomics analysis of malignant ascites from gastric cancer patients identified POSTN, a secreted protein, as one of four protein candidate markers of gastric cancer [53]. While not significantly more abundant in SKOV-3 spheroids incubated in Ascites 1, POSTN was significantly more abundant in Ascites 2-cultured spheroids compared to those formed in standard culture medium. Given the unique ability for Ascites 2 to drive spheroid aggregation and condensation, and the fact that POSTN expression has been linked to ovarian tumor invasion, dissemination [54], and recurrence [55], we hypothesized that the addition of POSTN to SKOV-3 cells grown in 3D, alongside chemokines also known to create a feedback loop that enhances cancer cell proliferation and invasiveness [56], would alter spheroid formation. We instead observed that the separate addition of these factors did not induce morphological changes that were observable through brightfield microscopy across a typical growth timeframe. Given the complex role of these factors *in vivo*, further studies could assess whether their addition may alter signaling pathways at the transcriptomic or proteomic levels.

SERPINA1, a member of the serine proteinase inhibitor protein family, was also uniquely detected in the spheroid-promoting Ascites 2 but absent in Ascites 1. However, it was also detected in SKOV-3 spheroids cultured in ascites, implying that incubation with ascites can induce SERPIN1 expression within cancer cell lines. Interestingly, our studies observed that SERPIN A protein abundances are significantly higher in spheroids collected directly from the ascites of chemotherapy sensitive patients, while SERPIN B proteins are significantly more abundant in those from resistant patients [29]. SERPINA10 has also been associated with improved progression-free and overall survival in the publicly available TCGA cohort, which was supported by further IHC-based validation in a patient cohort using fresh-frozen tissue lysates [57]. In opposition, a study in patient serum significantly associated SERPINA1 expression with platinum resistance [58].

Previous analysis of the effects of ovarian cancer ascites on adherent SKOV-3 cells visualized 300 protein spots on a two-dimensional gel and identified 20 proteins spots that were different. Upregulated proteins identified are PDZ domain-containing protein, GTP-binding nuclear protein Ran, zinc finger protein 268, collagen alpha 6(VI) chain, actin B, keratin type II cytoskeletal 8, Synaptotagami-like protein 3 and vimentin [59]; however, none were differentially abundant in our analyses of 2D or 3D cells cultured in ascites.

To ensure that the differences in the proteomes observed have not been caused by technical limitations, such as ascites derived proteins sticking nonspecifically to vessels and cells, we performed proteomic analyses of adherent SKOV-3 cells collected after washing. Data analysis revealed 22 secreted and ECM proteins that were upregulated in both spheroidal and adherent SKOV-3 cells cultured in ascites. Interestingly, TGM2 was the only secreted or ECM protein present in SKOV-3 cells grown in medium, which was further elevated when cells were cultured in ascites.

All other ascites-induced secreted or ECM proteins were only observed after ascites exposure. This pattern indicates that TGM2 expression is upregulated when cells are grown in ascites. Originally described as a transamidase protein that resides in the ECM, TGM2 is now known to be a GTP-binding protein that contributes to cancer progression through signal transduction, ECM modulation, and apoptosis regulation [60]. More recently, TGM2-deficient models have implicated this protein in spheroid formation. For instance, fewer and smaller spheroids coincided with elevated apoptosis in a fibroblast cell line [61], which has been reported to occur through reduced MEK1/2 and ERK1/2 activity [62]. Most recently, both knockdown and inhibition of TGM2 reduced spheroid formation in an epidermal squamous cell carcinoma model [63]. A new analysis of the GSE45553 public scRNA-seq dataset generated in a previous study [64] and downloaded from the GEO2R tool revealed TGM2 upregulation in lab-generated cisplatin-resistant OVCAR-8C spheroids versus their spheroids formed from their parental OVCAR-8 counterpart, supporting a previous observation of reduced overall survival and increased recurrence in patient populations [65].

Finally, the widespread adoption of three-dimensional ovarian cancer models, such as lab-grown spheroids, has enabled the investigation of personalized medicine strategies through *ex vivo* dose-response testing. However, most studies perform *ex vivo* drug response testing in personalized medicine approached with standard cell culture media, which may not accurately recapitulate the micro-environment of malignant ascites. We therefore assessed the influence of both culture conditions on the morphology of spheroids generated in ultra-low attachment plates. Spheroids generated from common ovarian cancer cell lines in ascites fluid showed vastly different morphology to those generated in cell culture medium, evidenced by the generation of more distinct and compacted spheroids. The addition of patient ascites encouraging the aggregation of these spheroids supports observations that most cells in malignant ascites are aggregated as spheroids following routine paracentesis.

## 4. Conclusions

The present study demonstrates that malignant ascites can support the formation of spheroids in ovarian cancer cells. Distinct morphological changes are observed in both cell line and ascites-derived spheroids in the presence of ascites, including increased compactness and aggregation. These phenotypical changes are reflected at the proteome level, where the abundances of proteins such as collagens, laminins, and annexin A2 were altered. In addition, elevated levels of transglutaminase 2 (TGM2), a known key player in cancer’s ability to resist treatment and spread, has been identified in SKOV-3 cells grown in ascites as 2D or 3D, indicating a direct role of ascites in the regulation of key protein expression. Despite these changes, which include increased viability observed in cells grown in ascites, chemoresponse is minimally affected and can still be monitored.

## 5. Materials and Methods

### 4.1. Cell Culture

The COV-362 (RRID: CVCL_2420) serous ovarian cancer cell line and A2780 (RRID:CVCL_0134) ovarian endometroid cell line, SKOV cells (SK-OV-3 [SKOV-3; SKOV-3], HTB-77) were purchased from the American Type Culture Collection (ATCC, Manassas, VA, USA).

All cell lines were authenticated via a short tandem repeat (STR) DNA profile in October 2022. The SKOV-3 cells were grown in DMEM media (Sigma Aldrich, St. Louis, MO, USA), cultured with the addition of 10% fetal bovine serum (Bovogen Biologicals, East Keilor, VIC, Australia) supplemented with 1% penicillin/streptomycin (Sigma Aldrich, St. Louis, MO, USA) and 1% L-glutamine (Sigma Aldrich, St. Louis, MO, USA).

### 4.2. Ascites collection

Ascites was collected from patients treated at the Royal Adelaide Hospital (RAH; Adeliade, Australia), with ethics approvals provided by the RAH (#140201) and Central Adelaide Local Health Network Human Ethics Committee (#R20181215), with patient consent and approval.

### 4.3. Primary HGSOC Culture

Primary cells were isolated from the ascites of advanced stage HGSOC patients (n = 3). All primary cells were grown in Advanced RPMI 1640 medium (Life Technologies, Mulgrave, VIC, Australia) supplemented with 4 mM L-glutamine, 10% FBS (Sigma Aldrich, St. Louis, MO, USA), and antibiotics (100 U penicillin G, 100 µg/mL streptomycin sulphate, and 100 µg/mL amphotericin B, Sigma Aldrich).

Cell aggregates were generated from primary ascitic ovarian cancer cells in plasma-treated flat-bottom plates and poly-HEMA–coated opaque ultra-low attachment plates. Primary cells were cultured in 0%, 1%, 5%, 10%, 50%, 75%, and 100% patient-matched ascites fluid mixed with advanced RPMI medium supplemented with 10% FBS, 1% penicillin/streptomycin (Sigma Aldrich, St. Louis, MO, USA) and 1% L-glutamine (Sigma Aldrich, St. Louis, MO, USA). Clinicopathological details are found in Supplementary Table 2.

### 4.4 Proteomics sample preparation

For serum and ascites samples, high-abundance proteins were depleted using the High-Select™ Top 14 Abundant Protein Depletion Spin Columns (Cat Number A36370, Thermo Fisher Scientific), following the manufacturer’s instructions.

For cellular samples, cells were maintained at 60–80% confluency for 3 passages prior to plating into 10 cm culture dishes. To estimate total cell numbers at harvest, a parallel dish was seeded with the same number of cells and counted manually. Following aspiration of the media, adherent cells were washed three times with 3 mL of pre-warmed PBS. Cell pellets containing up to 1 × 10^7^ cells were collected and washed three times with cold PBS, then stored at −80 °C prior to lysis.

For lysis, pellets were resuspended on ice in 200 µL RIPA buffer (Cat Number: 20–188, Millipore) supplemented with 1% (*v*/*v*) protease inhibitor cocktail (Cat Number: P8340, Sigma Aldrich). The lysate was sheared mechanically by passing it five times through a 26.5 G needle (Terumo, Cat. No. NN+2613R), then clarified by centrifugation at 20,000 × g for 30 minutes at 4 °C. The resulting supernatant was carefully transferred to a fresh tube.

Proteins were precipitated overnight at −20 °C by adding four-fold ice-cold acetone (Chem-Supply). The protein pellets were collected after centrifugation at 20,000g for 10 minutes at −9 °C, and the supernatant was discarded. Two additional washes with ice-cold acetone were performed, followed by centrifugation and removal of residual solvent. Pellets were air-dried on ice for 10 minutes before proceeding to downstream proteomic workflows.

Sample preparation for mass spectrometry analysis, followed the protocol by Doellinger *et al*. [66]. Briefly the cell pellets were resuspended in trifluoroacetic acid (TFA, Sigma-Aldrich). The mixture was incubated at room temperature for 2 minutes. The acidified samples were then neutralized with 2 M Tris base (Sigma-Aldrich, 10708976001) using 10 times the volume of TFA used for lysis. Reduction and alkylation were carried out by incubating the samples at 95°C for 5 minutes with Tris(2-carboxyethyl) phosphine (Merck) to a final concentration of 10 mM and 2-Chloroacetamide (Sigma-Aldrich) to a final concentration of 15 mM. After cooling to room temperature, the protein quantification was performed using tryptophan fluorescence [67]. Approximately 100 µg of the proteins from each sample were digested for 20 hours at 37°C with Trypsin Gold (Promega) at a protein-to-enzyme ratio of 1:100 ng. The digestion was stopped using TFA, and the samples were centrifuged at 20,000 g for 10 minutes. Peptides were then desalted using Sep-Pak C18 1 cc cartridges (Waters) and vacuum-dried at 50°C. The dried samples were resuspended in 10 µL of water, and the peptide concentration was estimated using a NanoDrop (Thermo Scientific, ND-ONE) at 205 nm. Before LC-MS analysis, samples were adjusted to a peptide concentration of 250 ng/µL and a final concentration of 0.1% formic acid (Merck).

### 4.5 Proteomics data acquisition

LC-MS analysis was performed on an EASY-nLC 1200 system (Thermo Scientific) coupled to an Orbitrap Exploris 480 mass spectrometer (Thermo Scientific). Approximately 1 µg of peptide sample was loaded at a flow rate of 600 nL/min onto an in-house packed C18 fused silica column (75 µm inner diameter, 360 µm outer diameter, 25 cm length) packed with 1.9 µm ReproSil-Pur 120 C18-AQ particles (Dr. Maisch). The column was heated at 50 °C and peptides were separated using a 70 min linear gradient of 3–20% acetonitrile in 0.1% formic acid at 300 nL/min. Two compensation voltages (−50 and −70 V) were alternately applied using a FAIMS Pro interface (Thermo Scientific) to adjust the entry of ionized peptides into the mass spectrometer. The mass spectrometer was operated in a data-dependent acquisition manner in positive ion mode. MS scans (m/z 300–1500) were recorded at resolution 60,000 (m/z 200). MS/MS scans were measured at resolution 15,000 after multiply charged peptide precursors (minimum intensity 10,000) were sequentially fragmented with higher energy collision dissociation (HCD) at 27.5% normalized collision energy. A dynamic exclusion period of 40 s was specified, and cycle times were restricted to 1.5 s.

### 4.6. Proteomics data processing

Raw FAIMS files were processed using FreeStyle 1.8 SP2 software (ThermoFisher Scientific), where data corresponding to individual CV settings (-50 and -70) were exported as separate raw files and subsequently analyzed using MaxQuant software version 1.6.3.4 [68]. The exported raw files from the two CV settings were treated as distinct fractions from the same sample. MS2 spectra were searched against the canonical reviewed Homo sapiens UniProt database (downloaded on 27^th^ September 2020). Enzyme specificity was set to “Trypsin/P”, the minimal peptide length was set to 7, and the maximum number of missed cleavages was set to 2. A maximum of five modifications per peptide were allowed; the mass tolerance for the first and main search of precursor ions was set to 20 ppm and 5 ppm, respectively, and the mass tolerance for secondary fragment ions was set to 0.02 Da. Carbamidomethylation of cysteine was searched as a fixed modification. “Acetyl (Protein N-term)” and “Oxidation (M)” were set as variable modifications. “Match between runs” and LFQ were activated. Results were filtered at a false discovery rate of 1% at the protein and peptide spectrum match level.

### 4.7. Proteomics data analyses

Reverse proteins, potential contaminants listed in the cRAP database [https://www.thegpm.org/crap/], and proteins inferred from peptides observed only at the MS1 level were excluded from subsequent analyses using Microsoft Excel. Tables were generated using Microsoft Excel, listing proteins only observed in all replicates of each condition.

Principal component analyses were performed and visualized using the EnhancedVolcano R package [69] and upset plots were generated using the UpsetR package [32]. The five most significantly enriched biological process in proteins uniquely expressed in Ascites 1 were identified using DAVID (NIH) [70] and visualized using a custom R-script that makes use of the ggplot2 package [71]. Bar plots were generated using GraphPad Prism 10 [www.graphpad.com].

### 4.8. Public molecular expression database

Single-cell RNA-seq data were exported from the GSE45553 dataset using the GEO2R online tool [72], which compared spheroids generated from the parental OVCAR8 (n=4) and cisplatin-resistant OVCAR-8C (n=4) cell lines. Data were originally obtained with the Affymetrix Human Gene 1.0 ST Array, and genes were considered significantly different in the context of a 1.5-fold or greater change in expression and a Benjamini & Hochberg-adjusted p-value lower than 0.05.

## Supporting information

All suppementary materials

## Supplementary Materials

Supplementary Figure S1: Primary ovarian cancer cells cultured as three-dimensional spheroids using plasma-treated flat-bottom and poly-HEMA–coated ultra-low attachment plates; Supplementary Figure S2: Heatmap showing protein abundance in top-14-depleted Ascites 1, 2, 3, and healthy serum; Supplementary Table S1: Alphabetized secreted, extracellular, and membrane proteins that were upregulated in SKOV-3 spheroids cultured in ascites compared to medium; Supplementary Table S2: Clinicopathological details of patients from which ascitic fluid and cells were sourced.

## Author Contributions: Conceptualization

M.K.H., P.H., M.K.O.; methodology: J.W., P.M., C.Y., M.K.H., P.H., J.S.; formal analysis: J.W., P.M., S.J., M.K.O, P.H., J.S.; resources, P.H.; M.K.O., M.K.H.; data curation: M.K.H., P.M., J.S.; writing: original draft preparation, M.K.H., M.P., J.S., writing—review and editing, M.P., P.H., M.K.O., C.Y., J.W., J.S.; supervision: M.K.O, P.H., P.M.; project administration: M.K.H, funding acquisition, M.K.H., P.H., M.K.O. All authors have read and agreed to the published version of the manuscript.

## Institutional Review Board Statement

The study was conducted according to the guidelines of the Declaration of Helsinki and approved by the Human Research Ethics Committee of Central Adelaide Local Health Network (CALHN). (HREC/18/CALHN/811, approved 14/02/2019).

## Informed Consent Statement

Informed consent was obtained from all subjects involved in the study.

## Data Availability Statement

The mass spectrometry proteomics data have been deposited to the PRIDE Archive via the PRIDE partner repository with the data set identifier PXD074417.

## Acknowledgments

The authors thank Dr. Noor A. Lokman for providing technical assistance, and Dr. Daniel Pincher for assisting with sample preparation.

