## Supplementary material for "Ascites-Driven Modulation of Cell Phenotypes and Proteomes: Implications for Cancer Progression": All suppementary materials

(b)

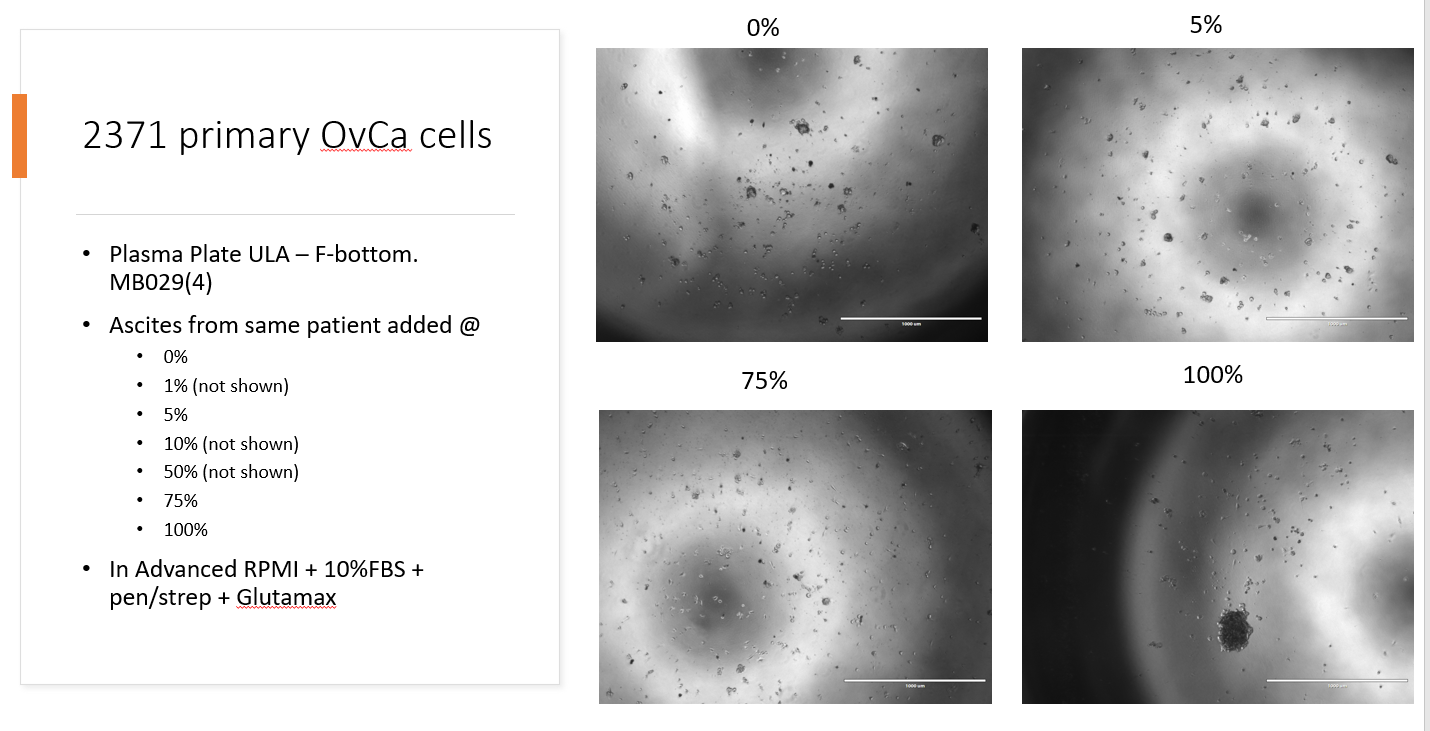
**(**a)

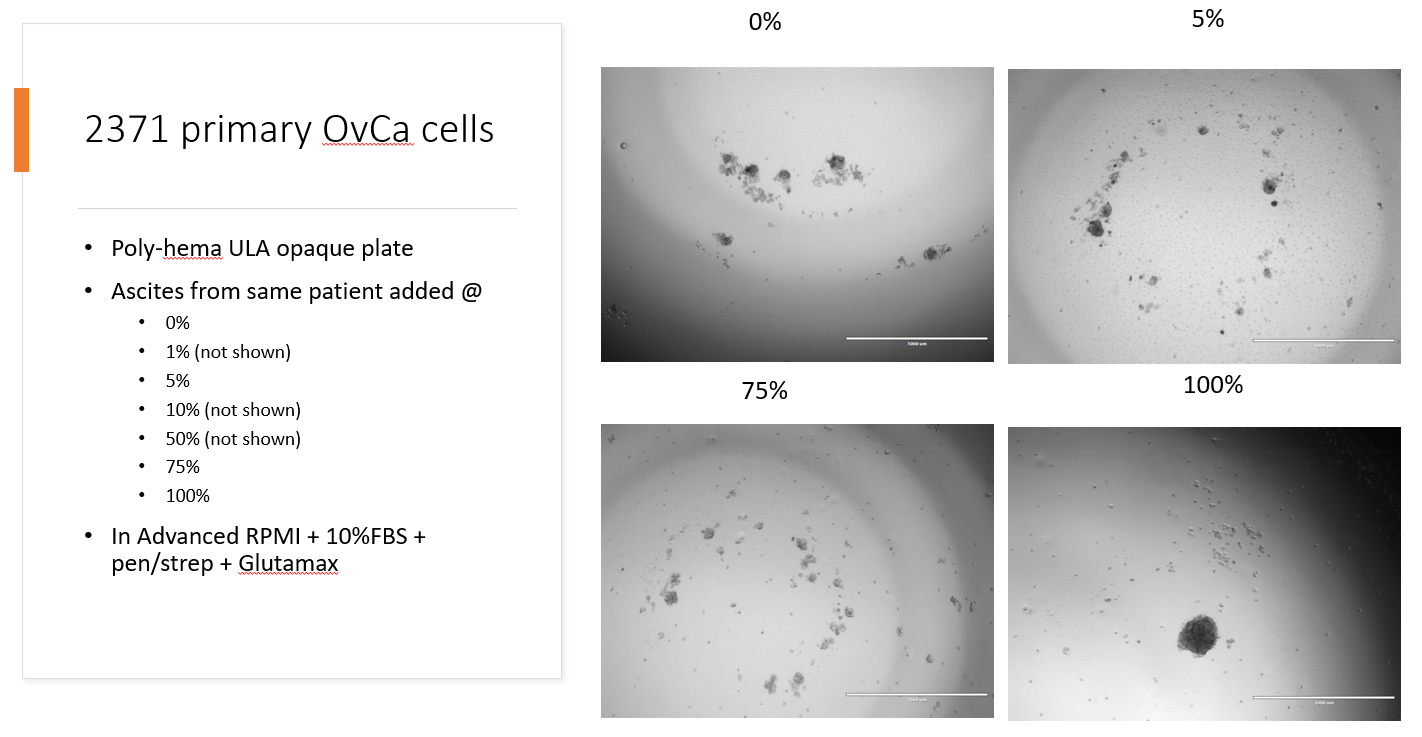

**Supplementary Figure S1:** Primary ovarian cancer cells cultured as three-dimensional spheroids using (a) plasma-treated flat-bottom and (b) poly-HEMA–coated ultra-low attachment plates at increasing concentrations of ascitic fluid in Advanced RPMI containing 10% FBS, 1X Penicillin-Streptomycin, and 2 nM L-glutamine.

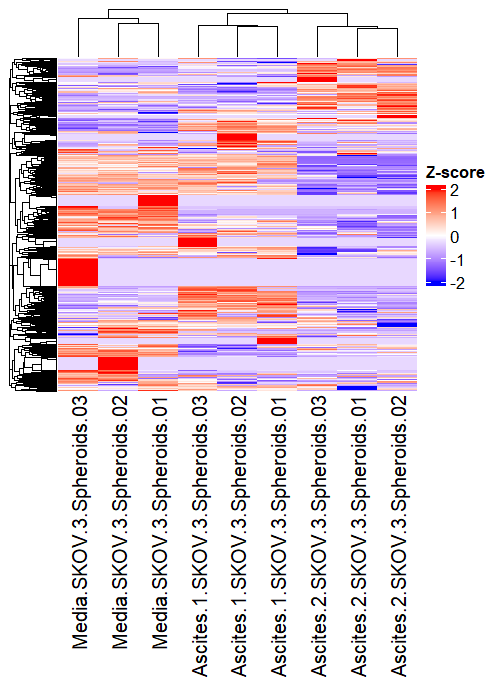

**Supplementary Figure S2:** Heatmap showing protein abundance in SKOV-3 cells grown in DMEM medium, Ascites 1 and Ascites 2 as 3D tumor spheroids. Protein abundance values are normalized and hierarchically clustered to highlight differences in expression patterns across conditions. Generated using the ComplexHeatmap R package.

**Supplementary Table S1:** Alphabetized secreted, extracellular, and membrane proteins that were upregulated in SKOV-3 spheroids cultured in ascites compared to medium. Relative abundances are an average of data from triplicate injections of spheroids cultured in ascites from two separate patients, or medium.

| **Gene** | **Description** | **Subcellular location** | **Ascites** | **Medium** |
| --- | --- | --- | --- | --- |
| A1BG | Alpha-1B-glycoprotein | Secreted | 8.39E+09 | Not detected |
| APOA1 | Apolipoprotein A-I | Secreted | 1.10E+11 | Not detected |
| APOA4 | Apolipoprotein A-IV | Secreted | 1.72E+10 | Not detected |
| APOC4 | Apolipoprotein C-IV | Secreted | 45317333 | Not detected |
| APOD | Apolipoprotein D | Secreted | 5.88E+09 | Not detected |
| C4BPB | C4b-binding protein beta chain | Secreted | 5.77E+08 | Not detected |
| COL1A2 | Collagen alpha-2(I) chain | Secreted, ECM | 1.55E+09 | Not detected |
| COL3A1 | Collagen alpha-1(III) chain | Secreted, ECM | 2.26E+08 | Not detected |
| COL5A1 | Collagen alpha-1(V) chain | Secreted, ECM | 27407667 | Not detected |
| COL5A2 | Collagen alpha-2(V) chain | Secreted, ECM | 2454667 | Not detected |
| CPB2 | Carboxypeptidase B2 | Secreted | 15003833 | Not detected |
| F10 | Coagulation factor X | Secreted | 93751667 | Not detected |
| F11 | Coagulation factor XI | Secreted | 1.56E+08 | Not detected |
| F12 | Coagulation factor XII | Secreted | 6.90E+08 | Not detected |
| GP1BA | Glycocalicin | Membrane | 24873167 | Not detected |
| HP | Haptoglobin | Secreted | 2.16E+11 | Not detected |
| HPX | Hemopexin | Secreted | 3.21E+10 | Not detected |
| ICAM1 | Intercellular adhesion molecule 1 | Membrane | 1.21E+08 | Not detected |
| MASP2 | Mannan-binding lectin serine protease 2 | Secreted | 5186333 | Not detected |
| PROZ | Vitamin K-dependent protein Z | Secreted | 4640500 | Not detected |
| GC | Vitamin D-binding protein | Secreted | 3.02E+10 | 1.32E+08 |
| VTN | Vitronectin | Secreted, ECM | 7.85E+09 | 6.41E+07 |
| COL1A1 | Collagen alpha-1(I) chain | Secreted, ECM | 2.36E+09 | 2.04E+07 |
| APOB | Apolipoprotein B-100 | Secreted | 1.64E+10 | 1.47E+08 |
| FGB | Fibrinogen beta chain | Secreted | 6.95E+10 | 1.04E+09 |
| CP | Ceruloplasmin | Secreted | 1.71E+10 | 5.53E+08 |
| PLG | Plasminogen | Secreted | 1.28E+10 | 4.58E+08 |
| F2 | Prothrombin | Secreted, ECM | 7.81E+09 | 7.32E+08 |
| ITIH3 | Inter-alpha-trypsin inhibitor heavy chain H3 | Secreted | 6.62E+08 | 1.03E+08 |
| APOE | Apolipoprotein E | Secreted, ECM | 5.39E+09 | 1.11E+09 |
| BST1 | ADP-ribosyl cyclase/cyclic ADP-ribose hydrolase 2 | Membrane | 34142000 | 7.67E+06 |
| LOXL2 | Lysyl oxidase homolog 2 | Secreted, ECM | 79468500 | 2.37E+07 |
| TGM2 | Protein-glutamine gamma-glutamyltransferase 2 | Membrane | 5.09E+09 | 1.68E+09 |
| COL6A2 | Collagen alpha-2(VI) chain | Secreted, ECM | 2.82E+08 | 1.08E+08 |
| CDH6 | Cadherin-6 | Membrane | 6.83E+08 | 2.65E+08 |
| CST3 | Cystatin-C | Secreted | 7.84E+08 | 3.30E+08 |
| SDC4 | Syndecan-4 | Membrane | 1.29E+08 | 6.16E+07 |

**Supplementary Table S2:** Clinicopathological details of patients from which ascitic fluid and cells were sourced.

| Patient | Age at diagnosis | Stage at diagnosis | Diagnosis |
| --- | --- | --- | --- |
| A1 | 48 | 3c | Serous carcinoma of the peritoneum |
| A2 |  |  | Serous carcinoma of the ovary |
| A3 | 70 | 3b | Serous papillary carcinoma of the ovary |
